# Bacteriophage diet breadth is impacted by interactions between bacteria

**DOI:** 10.1101/2023.06.05.543762

**Authors:** Ave T. Bisesi, Wolfram Möbius, Carey Nadell, Eleanore G. Hansen, Steven D. Bowden, William R. Harcombe

## Abstract

Predators play a central role in shaping community structure, function, and stability. The degree to which bacteriophage predators (viruses that infect bacteria) evolve to be specialists with a single bacterial prey species versus generalists able to consume multiple types of prey has implications for their effect on microbial communities. The presence and abundance of multiple bacterial prey types can alter selection for phage generalists, but less is known about how interactions between prey shapes diet breadth in microbial systems. Using a phenomenological mathematical model of phage and bacterial populations, we find that the dominant phage strategy depends on prey ecology. Given a fitness cost for generalism, generalist predators maintain an advantage when prey species compete, while specialists dominate when prey are obligately engaged in cross-feeding interactions. We test these predictions in a synthetic microbial community with interacting strains of *Escherichia coli* and *Salmonella enterica* by competing a generalist T5-like phage able to infect both prey against P22*vir*, an *S. enterica*-specific phage. Our experimental data conform to our modeling expectations when prey are competing or obligately mutualistic, although our results suggest that the *in vitro* cost of generalism is caused by a biological mechanism not represented in our model. Our work demonstrates that interactions between bacteria play a role in shaping ecological selection on diet breadth in bacteriophage and emphasizes the diversity of ways in which fitness trade-offs can manifest.

## Introduction

Predators often impose top-down control of ecosystems, impacting species abundances, community structure, and community function [1, 2]. For example, in marine environments, lytic bacteriophages (phages), the viral predators of bacteria, are critical drivers of microbial populations and nutrient cycling, lysing up to 40% of phytoplankton biomass per day [3]. The diet breadth of predators – how many different prey species they can consume – is an important component of how top-down control shapes an environment (Figure 1A) [4, 5, 6, 7, 8, 9, 10, 94]. Specialist predators often drive limit cycles with their prey, while generalists are more likely to stabilize prey populations through the emergence of apparent mutualisms, in which the presence of one prey species reduces the burden of predation on the other [13, 14, 15, 16, 17, 57]. Diversity in specificity is widespread in microbial communities, where some phages are generalists that can prey upon bacterial species across multiple genera, while others specialize on a single serovar [11, 12]. Identifying the forces shaping predator diet breath would therefore have substantial consequences for our ability to predict the long-term dynamics of multitrophic microbial communities.

**Figure 1.**
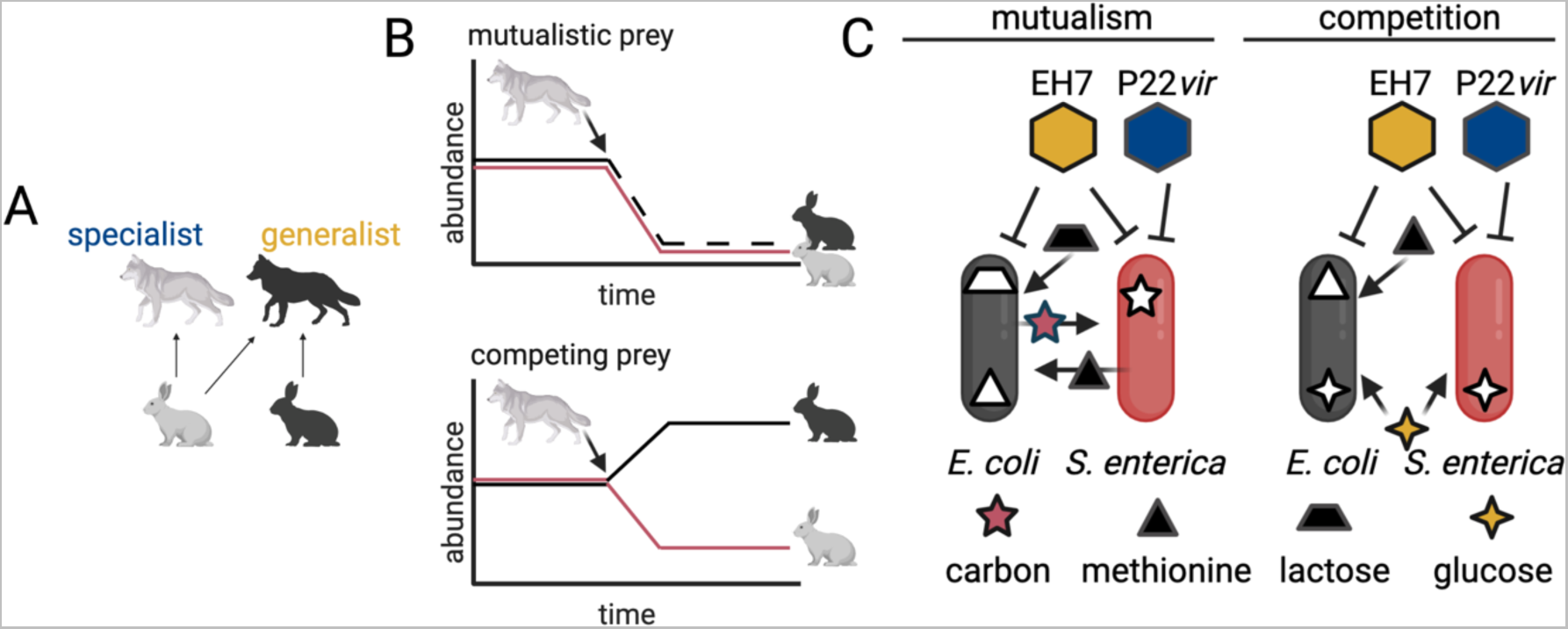
Diet breadth in a microbial synthetic community. **A:** Schematic of diet breadth. Species with generalist diets have wide diet breadths spanning multiple resources, while specialists have narrower diet breadths, sometimes specific to a single resource. **B**: Expected community dynamics when mutualistic or competing prey species are challenged by a specialist predator. When prey are mutualistic, predation will reduce abundances of both species. When prey compete, predation will reduce the abundance of one species and result in an increase in the abundance of the other species through competitive release. **C:** Schematic of cross-feeding system consisting of an *E. coli* methionine auxotroph and *S. enterica* methionine secreter. In lactose minimal media, *E. coli* provides carbon byproducts to *S. enterica* and *S. enterica* provides methionine to *E. coli*. In glucose minimal media with methionine, the two bacteria compete. The phage P22*vir* is a specialist on *S. enterica*, while the phage EH7 is a generalist that can attack both bacterial species. **For A, B, and C:** *Figure created with BioRender*.

The composition of the prey community is one force known to impact predator diet breadth. The evolution of generalist predators often requires prey heterogeneity to provide opportunities for diversification [18, 19, 20, 21, 22, 23, 34, 35, 100]. While it has been suggested that diversity in prey types could reduce the incidence of predator generalism, given the demands of engaging in coevolutionary arms races with more than one species [27], recent microbial studies have shown that the presence of multiple bacterial species was sufficient to select for generalists [18, 28]. Assuming a heterogenous prey environment, optimal foraging theory provides several additional predictions of how the structure of prey communities might shape predator diet breadth. First, it suggests that absolute prey densities alter selection on predator specificity by impacting foraging time. Generalism is predicted to be favored at low prey densities when foraging time is high, while specialism is favored at high prey densities; this prediction has been validated in a microbial system [26]. Optimal foraging theory also emphasizes the importance of relative prey abundances, such that predators should experience selection to exploit the most abundant prey types, even if those prey are low-quality or intraspecific competition between predators is strong [25, 32, 34, 35]. However, even when relative abundances are considered, most studies on diet breadth assume a static ratio of prey types over time, omitting a critical dimension of natural communities.

In particular, interactions between prey complicate the assumption of static ratios by generating correlations between prey abundances [29, 30, 31]. Competition for nutrients between bacteria tends to generate anti-correlated abundances between species, while positive interactions such as obligate mutualism tend to generate positively correlated abundances [29, 30, 31, 32]. When predators are consuming prey species with anti-correlated abundances, a generalist strategy is likely to be favored, because predation on one species should lead to an increase in the abundance of the alternative prey through competitive release (Figure 1B). Theoretical work provides some support for the notion that there are situations in which competing prey should favor generalist predators [32, 33]. When prey compete, mathematical modeling suggests that competitive dominance by a novel prey type is generally required for broadened predator diet breadth when fitness trade-offs for generalism are present [32]. Comparatively little work has been done investigating the impact of prey engaged in direct positive interactions on predator diet breadth. Positively correlated prey abundances are likely to favor specialist predators because predation by a specialist would also lead to a reduction in abundance of the alternative prey (Figure 1B). The interdependence between prey species may also increase the likelihood of overexploitation by predators [37, 38, 85]. However, these hypotheses - that competition between prey should favor predator generalism and that obligate mutualism between prey should favor predator specialism - have not been validated empirically.

Here we use a mathematical model and an *in vitro* system to investigate how interactions between prey species govern ecological selection on predator diet breadth. Our Lotka-Volterra type model incorporates two bacterial prey which either compete or engage in obligate mutualism, along with two phage predators, a specialist and a generalist (Figure 1C). The experimental system upon which our model is based is composed of *Escherichia coli* and *Salmonella enterica*, as well as two types of obligately lytic phage, a specialist and a generalist (Figure 1C). Our *S. enterica* strain secretes methionine and the *E. coli* strain used here is a methionine auxotroph [42]. These bacteria form an obligate mutualism in lactose minimal media, where *S. enterica* provides methionine to *E. coli* and *E. coli* secretes carbon byproducts that can be used by *S. enterica*. They compete in glucose minimal media when exogenous methionine is added to the media, because both species can use glucose and *E. coli* is not limited by methionine. P22*vir* is an obligately lytic phage specific to *S. enterica*. EH7 is an obligately lytic phage able to infect both *S. enterica* and *E. coli*.

We found that, in our model, obligately mutualistic interactions between microbial prey were more likely to favor a specialist phage predator, while competition between prey was more likely to favor a generalist phage predator. These findings were in accordance with our initial hypotheses and are broadly relevant to systems where interactions between prey drive correlations in their abundance. Our experiments recapitulated our ecological modeling results, although the biological mechanisms underpinning our observations were different than those considered in our model. Our work provides insight into how interactions between microbial prey species alter ecological selection on phage predator diet breadth and provides a foundation for predicting the evolution and maintenance of diet breath in bacteriophage, with implications for designing and managing microbial communities.

## Materials and Methods

### Model description

We constructed a model of the concentrations of two interacting bacterial species, a generalist phage, and a specialist phage. Bacteria (dimensionless biomass denoted by E or S) can either compete for resources or engage in obligate mutualism [adapted from Hoek et al., 2016; 44]. We assumed that prey interactions follow Lotka-Volterra-like dynamics, defining *μ_i_* as the maximum intrinsic growth rate of prey species *i*. We also considered a generalist predator (G) that could consume both prey species and a specialist predator (P) that could consume only one. Biomass of prey changes through growth (depending on the interaction with other prey species) and decreases due to predation and death/dilution:

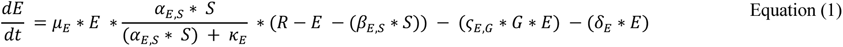

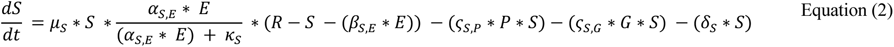

Biomass of predators increases through predation and decreases through death/dilution:

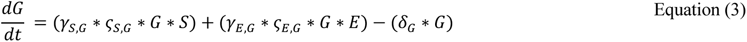

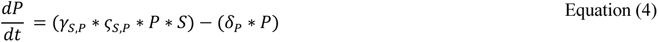

Our model was constructed such that three parameters determine prey interactions. Mutualism is determined by the saturation constant *κ_i_* and the mutualism coefficient *α_j_*_,*i*_, which reflects the beneficial effect of prey species *i* on the per capita growth of prey species *j*. If all *κ_i_* and *α_j_*_,*i*_ values are positive, bacterial species grow faster together and cannot grow alone. Competition is driven by the coefficient *β_j_*_,*i*_, where *β_j_*_,*i*_ determines the competitive effect of prey species *i* on the per capita growth rate of prey species *j*. Phage reproduction is modeled via adsorption with attachment rate *ς* which directly leads to lysis with burst size *γ*. The default natural death rate *δ_i_* was initially identical for all four species, as in a chemostat (Table 1).

**Table 1.** Phenomenological model parameters, default values, and descriptions.

| Parameter | Competition<br>Default Value | Mutualism<br>Default Value | Description |
| --- | --- | --- | --- |
| $\alpha_{j,i}$ | 1 | 1 | Mutualistic coefficient, benefit of prey species $i$ to prey species $j$ |
| $\beta_{j,i}$ | 0.9 | 0 | Competition coefficient, effect of prey species $i$ on prey species $j$ |
| $\mu_i$ | 0.5 | 0.5 | Intrinsic growth rate of prey species $i$ |
| $\gamma_{i,x}$ | 20 | 20 | Burst size of phage $x$ on prey species $i$ |
| $\varsigma_{i,x}$ | 0.001 | 0.001 | Attachment rate of phage $x$ on prey species $i$ |
| $\delta_i$ | 0.03 | 0.03 | Intrinsic death rate of species $i$ |
| $\kappa_i$ | 0 | 1 | Half-saturation constant of species $i$ |
| $R$ | 2 | 1 | System carrying capacity |

Using these equations, we investigated the extremes of pure mutualism (*κ_i_* and *α_j_*_,*i*_ > 0, *β_j_*_,*i*_ = 0), and pure competition (*κ_i_* = 0, *α_j_*_,*i*_ = 1, *β_j_*_,*i*_ > 0) (Table 1). *R* defined the carrying capacity of the system in each interaction type. The value of *R* was increased when prey competed relative to the value used when prey were mutualistic so that both phage types could be maintained across at least some parameter domains in each ecological condition.

### Model analyses

To compare how obligate mutualism and competition between prey affected relative abundance of the two predator types, we completed fixed point analysis to examine the fundamental behavior of the model and to determine the life history parameters of the predators expected to favor one phenotype over the other. Fixed point analysis of the model was performed in Mathematica 13.2.1.

We also investigated the equilibrium dynamics of the system by solving the ODEs until species abundances no longer changed between timepoints. All numerical simulations were completed in R v. 4.2.1 with the DeSolve package v. 1.32, using the LSODA solver with an absolute tolerance of 10^-14^. These results were verified by fixed point stability analysis. We evaluated equilibrium abundances of the phage predators under three different scenarios: 1) imposing a trade-off for expanded diet breadth by penalizing the burst size of the generalist phage, 2) altering the intrinsic growth rates or interaction coefficients of the bacterial prey, or 3) some combination of scenarios 1 and 2 (Table 2). To quantify phage coexistence, we used the equilibrium abundance of both phage. Relative abundance was calculated as the equilibrium density of the specialist divided by the sum of the equilibrium density of the specialist plus the equilibrium density of the generalist. Values greater than 0.5 indicated that the specialist was more abundant. Initial densities were the same across numerical simulations; all four species were always initialized at a density of 0.1. To confirm the significance of the parameters tested, we conducted two types of sensitivity analyses on our ODE system: the Morris screening method and the variance-based Sobol’ test [45, 46, 47, 48]. Morris screening and Sobol sensitivity analyses were performed in R with the ODESensitivity package v. 1.1.2 using the same parameter distribution ranges for each test type (Supplemental Table 3).

**Table 2.** Parameter trade-offs tested in our phenomenological model and their biological significance.

| Trade-off | Parameter Combinations | Significance |
| --- | --- | --- |
| None | $\gamma_{S,G} = \gamma_{S,P}$<br>and<br>$\varsigma_{S,G} = \varsigma_{S,P}$ | Generalist and specialist phage are parametrically identical |
| Cost of generalism | $\gamma_{S,G} < \gamma_{S,P}$<br>or<br>$\varsigma_{S,G} < \varsigma_{S,P}$ | Generalist and specialist phage differ in their abilities to kill prey |
| Interaction outcome | $\mu_i \neq \mu_j$<br>or<br>$\beta_{j,i} \neq \beta_{i,j}$<br>or<br>$\alpha_{j,i} \neq \alpha_{i,j}$ | Prey species coexistence in the absence of phage is biased or impossible |
| Cost of generalism and interaction outcome | $(\gamma_{S,G} < \gamma_{S,P} \text{ and } \mu_i \neq \mu_j)$<br>or<br>$(\gamma_{S,G} < \gamma_{S,P} \text{ and } \beta_{j,i} \neq \beta_{i,j})$<br>or<br>$(\gamma_{S,G} < \gamma_{S,P} \text{ and } \alpha_{j,i} \neq \alpha_{i,j})$<br>or<br>$(\zeta_{S,G} < \zeta_{S,P} \text{ and } \mu_i \neq \mu_j)$<br>or<br>$(\zeta_{S,G} < \zeta_{S,P} \text{ and } \beta_{j,i} \neq \beta_{i,j})$<br>or<br>$(\zeta_{S,G} < \zeta_{S,P} \text{ and } \alpha_{j,i} \neq \alpha_{i,j})$ | Generalist and specialist phage differ in their ability to kill prey and prey species coexistence in the absence of phage is biased or impossible |

Finally, following the *in vitro* finding of high rates of degradation of the generalist phage, we amended our model to impose a cost of generalism by increasing the death rate of the generalist relative to the three other species.

### Bacterial co-culture system and phage strains

The bacterial strains have been previously described [42]. Strains are listed in Supplemental Table 1. The *Salmonella enterica* serovar Typhimurium LT2 strain secretes methionine as a result of mutations in *metA* and *metJ* [51]. The *Escherichia coli* is a methionine auxotroph due to a deletion of *metB* [42]. To track bacterial abundances and relative ratios during growth, *E. coli* was tagged with a cyan fluorescent protein and *S. enterica* was tagged with a yellow fluorescent protein [52].

The specialist phage used was P22*vir*. It is an obligately lytic version of the lysogenic *S. enterica*-specific phage P22, created through several point mutations in its prophage repressor gene (Supplemental Table 6). P22*vir* was provided by I.J. Molineux. A generalist phage strain, EH7, was isolated and provided by E. Hansen and S. Bowden. EH7 is an obligately lytic T5-like siphovirus that uses BtuB, a differentially expressed outer membrane protein for vitamin B12 uptake, as a receptor. It is similar to T5-like coliphages described in Kim and Ryu (2011) [12] and Switt et al (2015) [53] (Supplemental Table 7).

Two additional bacterial strains were used for plaque assays (Supplemental Table 1). They were chosen so that, in mixed cultures of phage, phage types could be quantified independently of each other. The *E. coli* K-12 BW25113 Δ*trxA* from the Keio collection was used to quantify EH7 densities [96]. An *S. enterica* serovar Typhimurium NCTC 74 strain with *btuB* knocked out through a transposon insertion (EZ-Tn5 <KAN-2>, Lucigen) was used to quantify P22*vir*. The Δ*btuB S. enterica* strain was provided by S. Bowden.

### Media

Minimal hypho liquid media for experiments was prepared as previously described, with each component sterilized prior to mixing [99] (Supplemental Table 2). In addition to the appropriate carbon source, solutions containing sulfur, nitrogen, phosphorus, and metals were supplemented into each media type (Supplemental Table 2). Routine culturing of all bacterial strains was carried out on Miller Lysogeny Broth (LB) unless otherwise indicated. Working stocks of both phage types were grown on log-phase *S. enterica* LT2 cultures in LB and stored at 4°C. Stock titer was determined by plaque overlay assay on the appropriate strains as described above.

### Phage competition assays

Phage competition assays were performed in 96-well flat bottom plates on a Tecan Infinite Pro200 plate reader for 48 hours at 37°C with shaking at 432 rotations per minute. Experiment duration was chosen to allow batch culture experiments to reach a final state (stationary phase, phage densities unchanging), thus allowing us to compare to our chemostatic model. Overnight stationary phase cultures in LB started from single colonies were washed three times in saline, adjusted to a density of 10^7^ cells per mL, and used to inoculate 200µL of appropriate medium with 2.0 × 10^5^ total cells per well (i.e. 2.0 × 10^5^ total *S. enterica* cells in monoculture, 1.0 × 10^5^ total *S. enterica* cells and 1.0 × 10^5^ total *E. coli* cells in co-cultures). Phage stocks were diluted in saline to 10^5^ plaque-forming units per mL and inoculated into the appropriate wells to an MOI between 0.005 and 0.01, depending on the fraction of infectable cells for each phage type. Phage strains were added either in isolation (10^3^ total phage particles of either P22*vir* or EH7) or in a one-to-one ratio (10^3^ total phage particles of P22*vir* and 10^3^ total phage particles of EH7 for a final density of 2 × 10^3^ total phage particles). Phage densities were confirmed by plaque overlay assay at the start of the experiment on the appropriate strains as described below.

OD600, *E. coli*-specific CFP (Ex: 430 nm; Em: 480 nm), and *S. enterica*-specific YFP (Ex: 500 nm; Em: 530 nm) fluorescence were read every 20 minutes. Fluorescent protein signals were converted to species-specific OD equivalents using an experimentally determined conversion factor as previously described [54]. A single initial experiment was completed to confirm the reproductive ability of each phage on *S. enterica* monoculture or *E. coli* monoculture. Full factorial experiments testing all three phage conditions (P22*vir*, EH7 or EH7 + P22*vir*) on either *S. enterica* monoculture, mutualistic co-culture, or competitive co-culture were then completed with four biological replicates per condition, plus three biological replicates for no-phage controls per condition. Three experiments were set up and completed during different weeks to confirm the repeatability of the results; one representative run was chosen for display in this paper. We plated for PFUs for each replicate from half of the total 200µL volume using plaque overlay assays on LB plates with 0.7% LB top agar at the end of the 48 hour growth period. All replicates were quantified using plaque assays on both Δ*trxA E. coli* and Δ*btuB S. enterica.* Δ*trxA E. coli* and Δ*btuB S. enterica* were prepared for use in plaque assays through overnight culture growth in LB, prior to being diluted 1:10 (Δ*btuB S. enterica*) or 1:5 (Δ*trxA E. coli*) and allowed to grow for 30 minutes. Plaque assays were otherwise performed as previously described using 2µL of phage spot dilutions from 10^0^ to 10^-7^ with three technical replicates per dilution per sample [55, 56]. The lower limit of detection was 500 PFU/mL. Change in phage titer was represented as the natural log of the final phage density divided by the starting phage density (ln(final PFU/mL / initial PFU/mL)). All plates were incubated overnight at 37°C.

To impose a cost of generalism in our system, we repeated the phage competition assays as previously described, incubating the phage in minimal media at 37°C with shaking for 24 hours prior to the addition of cells in either *S. enterica* monoculture or competitive co-culture. Mutualistic co-culture was not tested. The experiment was completed once following preliminary trials to confirm that EH7 did not degrade below the limit of recovery after 24 hours. Once cells were added, cultures were grown for an additional 24 hours. Phage densities were quantified at the beginning and end of the 48 hour experiment.

### Phage degradation assays

We examined the impact of cell starvation on the formation of new EH7 particles using a full factorial design of both phage types and *E. coli* or *S. enterica* monoculture in lactose hypho minimal media. Neither bacterial strain could grow, as each was starved of essential nutrients. Bacteria were inoculated in lactose hypho monoculture at a density of 10^5^ cells per 200µL of medium in a 96-well plate and treated with either EH7 or P22*vir* at a total density of 10^3^ PFU (MOI = 0.01) or incubated in a no-phage control. Additionally, we tested each phage in isolation in lactose minimal media without cells to determine phage decay rates. Each condition consisted of three replicates. Both experiments were completed once at 37°C with shaking at 432 rotations per minute. Phage density in each well was determined by plaque overlay assay at the beginning and end of the 48 hour experiment. Change in phage titer was again represented as the natural log of the final phage density divided by the starting phage density (ln(final PFU/mL / initial PFU/mL)).

### Scripts and data availability

Numerical simulations, sensitivity analyses, data analysis, statistics, and figure generation were performed using R v. 4.2.1 using custom scripts available at https://github.com/bisesi/Host-Ecology-and-Host-Range. Raw experimental data and Mathematica notebooks for fixed point analysis are available at the same link.

## Results

### In a phenomenological model, phage relative abundance depends on prey interactions and fitness trade-offs

We used a phenomenological model to predict how communities of two interacting prey species respond to attack by predatory lytic phage during chemostatic growth. We predicted that competition between prey species was likely to favor predator generalism by increasing temporal heterogeneity in resource availability [36, 98], while obligate mutualism between prey species would result in less temporal heterogeneity, as bacterial species would either occur together or not at all, likely favoring specialization [36].

We first investigated the behavior of the model with a single parameter set. Using our default parameters (Table 1), we examined cases in which phage were not present, or when only one phage type was present. When phage were not modeled in our system, prey species reached a 50:50 ratio at equilibrium (Figure 2A, left panel). The introduction of a specialist resulted in competitive release of *E. coli* when prey competed and correlated reductions in bacterial abundances when prey were mutualistic. This behavior was consistent with our conceptual model (Figure 2A, middle panel; see also, Figure 1B). In comparison, the introduction of a generalist phage predator reduced the density of both prey species at equilibrium, keeping their abundances at a 50:50 ratio (Figure 2A, right panel). We also considered the behavior of the model when phage phenotypes competed against one another. In alignment with our conceptual model, when the specialist had a burst size five times greater than that of the generalist, the generalist dominated when prey competed (Figure 2B), while the specialist dominated when prey were mutualistic (Figure 2C).

**Figure 2.**
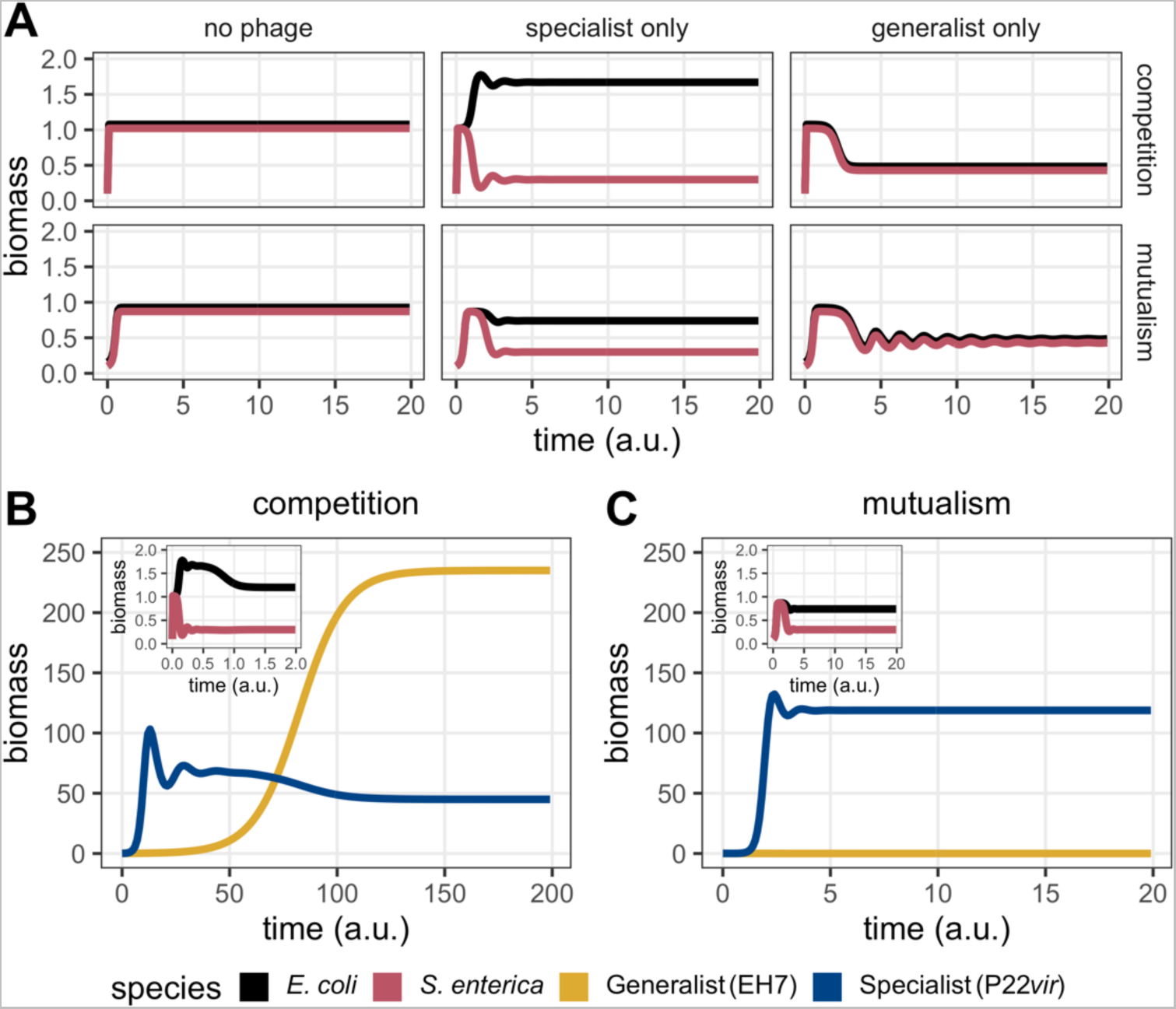
Numerically-simulated bacterial dynamics demonstrate that competing prey provide a different selective environment for phage than mutualistic prey. **A:** In the absence of phage, both prey species reach an equilibrium ratio of 50:50. In the presence of only a specialist phage, the prey attacked by the specialist (pink line, *S. enterica*) decreases. The prey that is not attacked (black line, *E. coli*) reaches a higher equilibrium frequency if the prey are competing and a lower equilibrium frequency if the prey are mutualists. When only the generalist predator is present, regardless of prey interactions, prey remain at a 50:50 ratio throughout the simulated growth period. **B:** When prey compete and both phage types are present, the generalist (yellow line, EH7) dominates over the specialist (blue line, P22*vir*). The fraction of the bacterial population consisting of *S. enterica* decreases significantly, while *E. coli* proliferates (inset plot). **C:** When prey are mutualistic and both phage types are present, the specialist (blue line, P22*vir*) dominates over the generalist (yellow line, EH7). The fraction of the bacterial population consisting of *S. enterica* decreases substantially less than in the case of competing prey, though the overall bacterial population is suppressed relative to the no-phage condition (inset plot). **For B and C:** System dynamics represent parameter space in which the specialist’s (blue line, P22*vir*) burst size is five times that of the generalist (yellow line, EH7).

To further understand the behavior of the model when phage phenotypes competed against one another, we then sought to simplify the system, focusing on Eqs 3–4. *R*\* theory [58] suggests that, when two species compete for the same resource, the species with the lower *R*\*-that is, the species that can sustain zero net population growth on fewer resources - will be able to competitively exclude the other species. To apply this to our system, we treated the shared prey, *S. enterica*, as a resource, and identified domains in which the amount of *S. enterica* required by the specialist to hold its net population growth at zero, 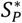, would be expected to be less than the amount of *S. enterica* required for zero net growth of the generalist population, 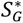 [58]. To do so, we assumed that the generalist phage had an equivalent burst size and attachment rate on both prey types and that the intrinsic mortality rate was not species-specific, such that *γ_G_* described the burst size of the generalist on both prey, *ς_G_* described the attachment rate of the generalist on both prey, *γ_P_* described the burst size of the specialist on *S. enterica*, *ς_P_* described the attachment rate of the specialist on *S. enterica*, and *δ* described the intrinsic mortality rate of all four species.

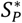 and 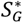 were obtained by setting the left-hand side of Eqs 3–4 equal to 0, and solving for *S*. Using these equations, in accordance with R* theory, requiring that

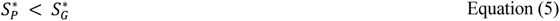

leads to:

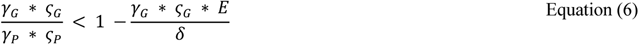

This analysis therefore suggested that the specialist phage could dominate (i.e. have the lower *S*\*) when the alternative prey source *E. coli* was rare, or when the generalist suffered a fitness trade-off for expanded diet breadth in the form of reduced burst size or attachment rate. This inequality is broadly consistent with the “jack of all trades, master of none” hypothesis for the predominance of specialization [24, 59, 74, 75, 76, 77, 78, 79, 92, 95, 97].

To verify the intuition of our *S*\* inequality, we used our default parameters (Table 1) to consider various domains in which the specialist phage predator was favored when prey species converged to a 50:50 ratio at equilibrium in the absence of predators. To do so, we systematically changed the burst size (Figure 3; for attachment rate *ς_S,P_*, see Supplemental Figure 1) of the specialist phage on the shared prey *S. enterica* (*γ_S,P_*) and considered the relative abundance of each phage type at equilibrium. We found that even as the cost of phage generalism increased, the generalist phage was always favored when prey competed (Figure 2B; Figure 3A, left panel). Fixed point analysis demonstrated that if *ς_P_* > 2*ς_G_* or *γ_P_* > 2*γ_G_*, i.e., specialist adsorption rate or burst size was twice as large as that of the generalist, then the only stable fixed point was a four-species equilibrium. No domain existed within relevant parameters where only the specialist phage could stably coexist with the two prey species. These results suggested that, even in those cases where the generalist was not dominant, it could not be driven extinct by a specialist when prey competed. Conversely, the specialist was favored on mutualistic prey (Figure 2C; Figure 3A, right panel) given a minimum cost of generalism. If *ς_P_* > 2.83*ς_G_* or *γ_P_* > 2.83*γ_G_*, i.e. the specialist’s burst size or attachment rate was 2.83 times as large as that of the generalist, then the only stable fixed point was that of the two bacterial species coexisting with the specialist. As such, given a threshold cost of generalism, only the specialist could coexist with mutualistic bacterial species, regardless of initial conditions.

**Figure 3.**
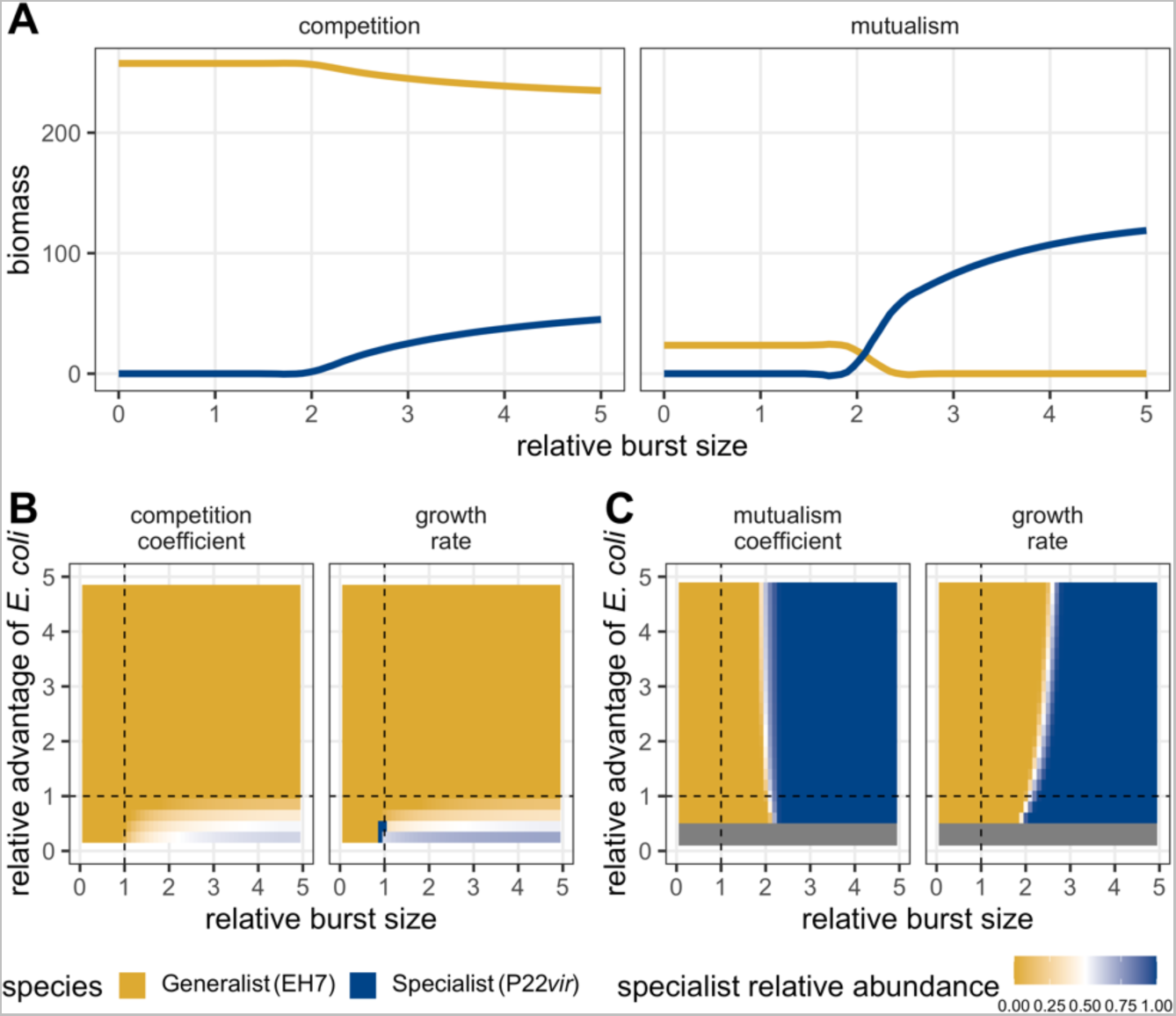
Numerically-simulated phage dynamics given a variety of parameter trade-offs demonstrate that prey interactions result in different patterns of predator abundance. **A:** The final density of each phage type as a function of bacterial interactions and increasing cost of generalism modeled as increasing specialist burst size. When prey are mutualistic, a cost of generalism exists that favors specialists (blue line, P22*vir*) over generalists (yellow line, EH7). When prey compete, no such cost exists; this is true even as the specialist’s burst size increases well beyond the values displayed here. **B:** The relative abundance of the specialist phage on competing prey as a function of increasing cost of generalism and relative growth advantage of the alternative prey *E. coli*. Whether prey growth advantage is modeled through growth rate or competitive coefficients (beta), the generalist is favored (yellow, EH7) except in a small subset of cases where the alternative prey is competitively excluded. **C:** The relative abundance of the specialist phage on mutualistic prey as a function of increasing cost of generalism and relative growth advantage of the alternative prey *E. coli*. Whether prey growth advantage is modeled through growth rate or mutualistic benefit (alpha), a cost of generalism exists above which specialism is favored (blue, P22*vir*). Note that there are benefit and growth rate values for *E. coli* below which the mutualistic system cannot be supported, indicated by the grey bar.

To visualize the next prediction of our *S*\* inequality - that the relative availability of the alternative prey source *E. coli* mattered for the competitive outcome of phage diet breadth strategies - we loosened the previous assumption that bacterial species should reach a 50:50 ratio without predators. In addition to varying the ratio of prey growth rates (in both competition and mutualism) or the ratio of interaction coefficients (*α_j_*_,*i*_ for mutualism, *β_j_*_,*i*_ for competition), we systematically altered the cost of generalism (as both burst size and attachment rate; for attachment rate, see Supplemental Figure 1). Our analyses demonstrated that when prey were competing, relative growth rates and competition coefficients mediated the abundance of the specialist such that it had a fitness equivalent to or greater than that of the generalist only in those cases where the alternative prey *E. coli* was competitively excluded by the shared prey *S. enterica* (Figure 3B). However, even in these cases, only the four-species equilibrium point was stable, suggesting that even when the specialist dominated due to the competitive superiority of its prey, the generalist was able to coexist. In contrast, when prey were mutualistic, neither relative growth rates nor relative mutualistic benefit coefficients altered phage relative abundance greatly. Instead, ecological dominance of the specialist depended mainly on the cost of generalism, such that the specialist tended to proliferate given a sufficient cost regardless of biased bacterial abundances driven by unequal mutualistic benefit (Figure 3C).

Finally, to ensure that our tested parameters captured the fundamental behavior of the model, we performed two sensitivity analyses - a Morris screening and a Sobol variance analysis - on our ODE equations to determine which parameters had the largest impact on the final biomass of each phage type. In the case of both obligate mutualism and competition, Morris screening methods suggested that the death/dilution rate, burst size and attachment rates of both phage, and the interaction parameters for the microbial species were of greatest impact (Supplemental Table 5). The variance-based Sobol method reinforced the importance of dilution rate (Supplemental Table 6). These results were consistent both with the parameters identified by fixed point analysis, our *S*\* inequality, and the basic construction of the model, which requires that both phage types have reproductive parameters sufficient to offset the chemostat-induced mortality rate.

### Phage relative abundance *in vitro* aligns with modeling results in co-culture

Using our wet-lab experimental cross-feeding system, we tested the mathematical prediction that generalist predators would be favored on competing prey and specialist predators would be favored on mutualistic prey. We first verified that over 48 hours our specialist phage (P22*vir*) could only replicate on *S. enterica* and that our generalist phage (EH7) could replicate on both prey species (Figure 4A). Each phage alone also grew well on competitive and mutualistic bacterial co-cultures. The final density of EH7 appeared similar when replicating on both interaction types (Figure 4A, p = 0.999). This was true for P22*vir*, as well, such that type of interaction did not result in significantly different final titers (Figure 4A, p = 0.909). However, when phage were added in isolation, EH7 reached a higher titer than P22*vir* on both competitive (Figure 4A, p < 0.0001) and mutualistic co-culture (Figure 4A, p < 0.0001). Consistent with our model, P22*vir* increased *E. coli* frequency relative to the no-phage control in competitive co-culture (Figure 4C, p < 0.0001). Applying the specialist phage also suppressed, but did not eliminate, both bacterial species in mutualism (Figure 4D). In contrast, the generalist phage more effectively suppressed *E. coli* in competitive co-culture, resulting in a higher relative density of *S. enterica* compared to the no-phage control (Figure 4C, p = 0.0035). Unlike P22*vir*, the application of EH7 to mutualistic co-culture did not suppress co-culture growth for the duration of the experiment (Figure 4C).

**Figure 4.**
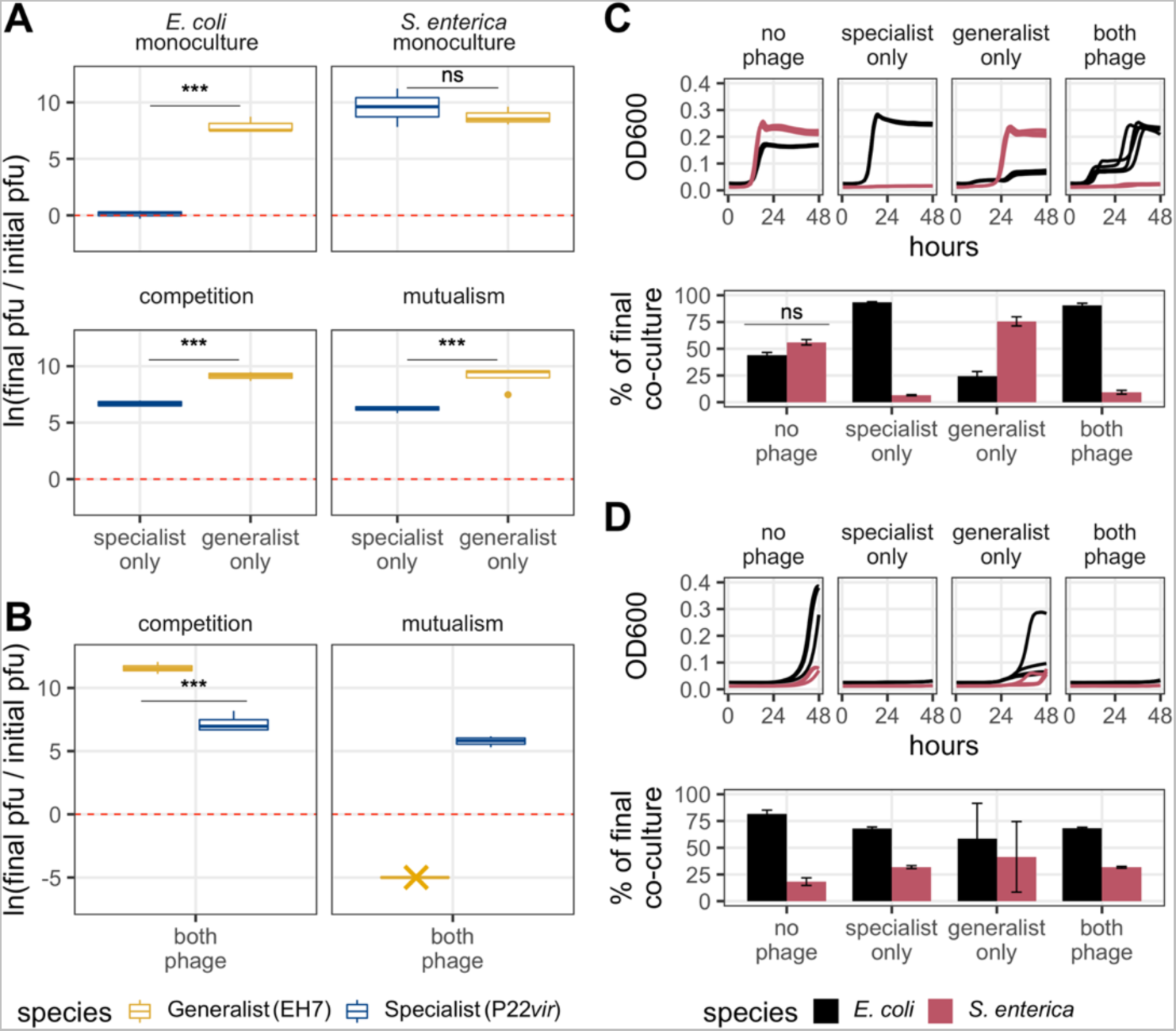
Phage and bacterial dynamics *in vitro* align with modeling expectations. **A:** Change in individual phage titer on bacterial monocultures and co-cultures over a 48 hour time period. P22*vir* replicates effectively on *S. enterica* but not *E. coli*, while EH7 grows on both bacterial strains. EH7 reaches approximately equivalent densities on both interaction types (p = 0.999), as does P22*vir* (p = 0.909), although its final density is reduced compared to EH7 (competition: p < 0.0001, mutualism: p < 0.0001). **B:** Change in phage titer when phage compete on different bacterial interaction types over a 48 hour time period. When competing against the specialist, EH7 dominates when prey compete (p < 0.0001), doubling more times than in isolation (p < 0.0001). P22*vir* dominates when prey are mutualistic (p < 0.0001), while EH7 disappears below the limit of detection, as indicated by the yellow asterisk. **For A and B:** The dotted red line indicates no change in titer from the start of the experiment to the end. Values greater than zero indicate an increase in titer, while values below zero indicate a decrease in titer. Statistical significance was determined using a two-way ANOVA with Tukey’s HSD multiple comparison test. **C:** Species-specific ODs over time across treatment conditions (top) and final fraction of bacterial co-culture composed of each strain across treatment conditions (bottom) when bacteria compete. The fraction of *E. coli* increases in those cases in which *S. enterica* is predominantly suppressed by phage (specialist only and both phage treatments). *E. coli* frequency is reduced relative to no-phage controls when only EH7 is applied (p = 0.0035). **D:** Species-specific ODs over time across treatment conditions (top) and final fraction of bacterial population composed of each strain across treatment conditions (bottom) when bacteria are mutualistic. *E. coli* dominates at similar levels in all treatment conditions, even in cases where overall growth is suppressed. **For C and D:** Statistical significance for bar graphs was determined using a two-way ANOVA with Tukey’s HSD multiple comparison test. OD600 traces represent 4 biological replicates for conditions with phage and 3 biological replicates for conditions without phage.

We also evaluated population dynamics when the two phages competed against one another (Figure 4B, 4C and 4D). When both phage were present in competitive co-culture, the generalist reached a higher final density than the specialist (Figure 4B, p < 0.0001), as predicted by our modeling results. Bacterial dynamics were also consistent with our model, with *E. coli* dominating through competitive release (Figure 4C, p < 0.0001). When both phage were present in mutualistic co-culture, the specialist reached a higher final titer (Figure 4B, p < 0.0001) and co-culture densities were suppressed for the duration of the experiment (Figure 4D). Curiously, however, the presence of the specialist phage reduced EH7 below the limit of detection when prey were mutualistic (Figure 4B), a result that we interrogated further.

### *In vitro*, cost of generalism manifests as increased rate of degradation and reduced infectivity of starved cells

To understand our inability to detect the generalist phage in mutualistic co-culture at the end of our experimental window, we investigated the ability of each phage to reproduce on starved cells by adding phage to monocultures in lactose minimal media as previously described (see Materials and Methods). With *E. coli* or when placed in wells without bacterial cells, P22*vir* titer remained unchanged over the 48 hour growth period (Figure 5A). When placed in wells with starved *S. enterica*, P22*vir* was able to reproduce despite the expected physiological inaccessibility of its prey in a nutrient-deprived state, with its titer increasing relative to the condition without cells (Figure 5A, p = 0.001) or with *E. coli* (Figure 5A, p = 0.0001).

**Figure 5.**
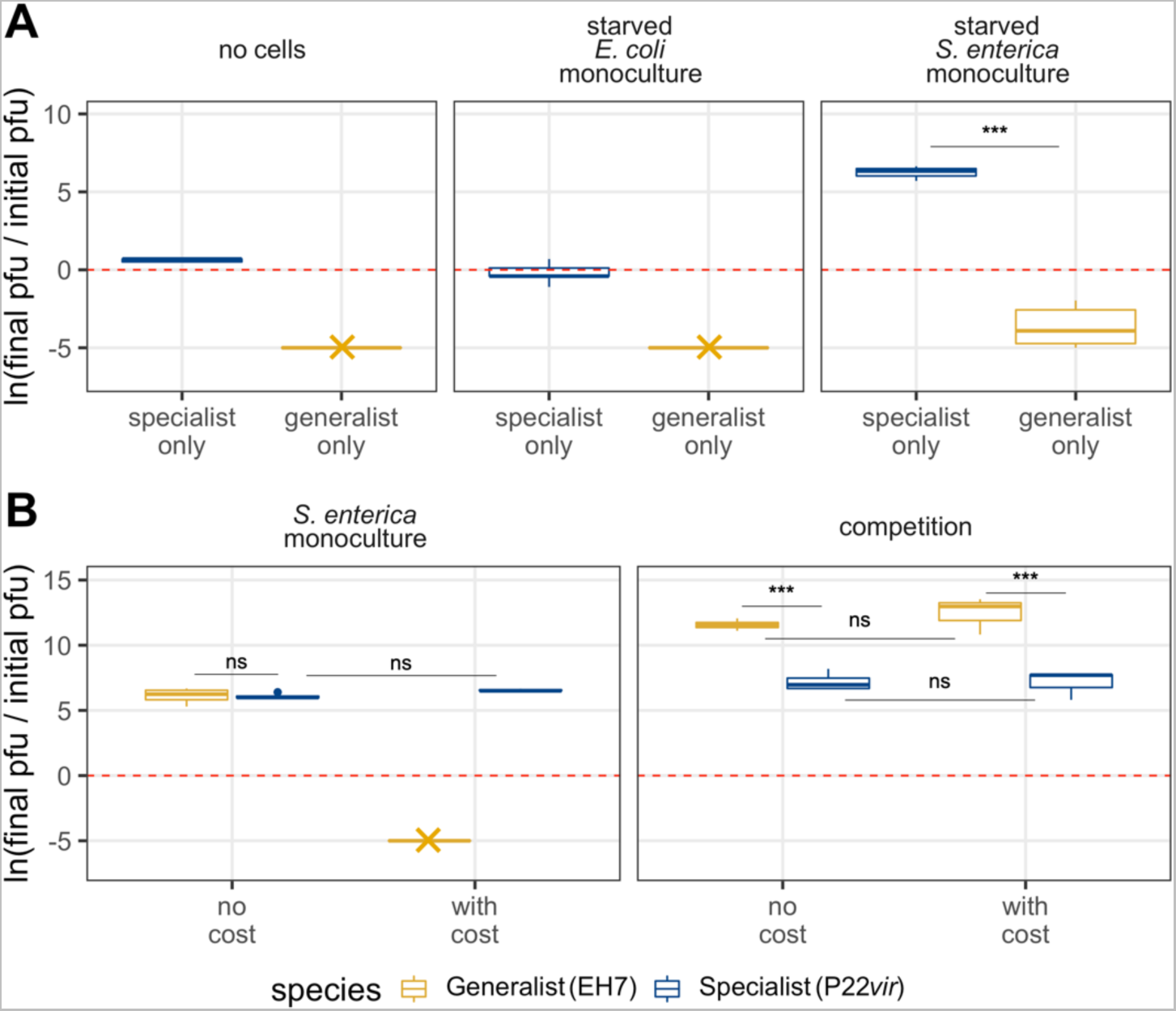
Imposing a cost of generalism *in vitro* as phage intrinsic mortality recapitulates modeling results on *S. enterica* monoculture. **A:** Change in phage titer across interaction types and phage treatments when cells are starved or not present in cultures. P22*vir* does not degrade over the 48 hour growth period, even in cases when there are no cells present that it can productively infect. Additionally, it can reproduce even on starved *S. enterica*. EH7 degrades below the limit of detection in all conditions, as indicated by the yellow asterisk, but does not disappear completely when *S. enterica* is present, suggesting *S. enterica* may be somewhat more physiologically accessible to it than *E. coli* in a starved state. **B:** Change in phage titer across interaction types when cells are either added to minimal media at the same time as phage (no cost) or 24 hours later (with cost). Results are shown for phage competition assays, when both phage types are present in wells at the start of the experiment. In the no cost condition, EH7 reaches comparable titer to P22*vir* on *S. enterica* monoculture (p = 0.999) and wins when prey compete (p < 0.0001). In the condition where cells are added after a period of phage incubation, EH7 degrades below the limit of detection on *S. enterica* monoculture but dominates when prey are competing (p < 0.0001). **For A and B:** The dotted red line indicates no change in titer from the start of the experiment to the end. Values greater than zero indicate an increase in titer, while values below zero indicate a decrease in titer. Statistical significance was determined using a two-way ANOVA with Tukey’s HSD multiple comparison test.

In comparison, the generalist phage EH7 decreased in abundance in all conditions without growing cells after the 48 hour growth period. There were no detectable phage particles in wells without cells or with starved *E. coli.* In wells with only starved *S. enterica*, some phage were still detectable, although phage titer was greatly reduced, suggesting that *S. enterica* may be more physiologically accessible to EH7 than *E. coli* in a starved state (Figure 5A, p = 0.012). Taken together, these results indicate that the generalist phage suffers a cost that manifests in two distinct ways: first, a rapid rate of degradation in minimal media, and, second, a reduced capacity to infect and reproduce inside starved cells relative to the specialist phage P22*vir*. As a result, when competing with P22*vir* on mutualistic co-culture, the starved physiology of the interdependent cells and the generalist’s degradation can explain our inability to detect it at the end of previous experiments.

Interestingly, these results are specific to minimal media, as EH7 does not degrade in LB (Supplemental Analysis 2; Supplemental Figure 2). We were not able to identify which component of our minimal media was responsible for the degradation of the phage, though it does not appear to be related to the presence of metals or the result of osmolarity (Supplemental Analysis 2; Supplemental Figure 2).

### The generalist is favored in competition even when a cost is imposed *in vitro*

While our *in vitro* experiments replicated our modeling results when bacterial prey were mutualistic or competing, we did not observe a cost of generalism when both phage types were competing for the shared prey *S. enterica*, with each phage reaching titers that did not differ significantly from one another (Figure 5B, p = 0.999). In our system, a trade-off may thus only manifest in the mutualistic treatment where the generalist phage degrades due to bacterial growth delayed by P22*vir* predation on *S. enterica*. To impose a cost of generalism across treatments, we repeated our phage competition assay experiments, incubating the phage for 24 hours prior to the addition of cells, anticipating that some degradation of the generalist EH7 would occur, while the titer of P22*vir* would remain unchanged. Previous preliminary experiments had indicated that while EH7 titer is often below the limit of detection after 24 hours (Supplemental Figure 2), the phage can be recovered following the addition of cells, suggesting that some infectious phage particles remain. We therefore expected that the reduced titer of EH7 when cells were applied would mirror the conditions of mutualism when phage competed. Given that the presence of P22*vir* in competitive co-culture increases *E. coli* frequency, we hypothesized that a lower titer of EH7 when cells are added may still result in a higher final titer of the generalist phage relative to the specialist, because any remaining EH7 would be able to utilize *E. coli*.

When a cost was imposed and phage were competed on *S. enterica monoculture*, the generalist disappeared below the limit of detection, while P22*vir* final titer was not significantly different than in the condition without cost (Figure 5B, p = 0.982). In contrast, we found that when bacteria competed, final titers of both EH7 (Figure 5B, p = 0.723) and P22*vir* (Figure 5B, p = 0.999) were comparable to the no cost condition, as we had expected. These results are consistent with our modeling results, suggesting that, even with a cost of generalism, a generalist predator should be favored on competing prey.

### In a phenomenological model, prey interactions determine the intrinsic death rate needed to favor specialism

Finally, we amended our model to see if we could replicate the results of our *in vitro* experiment, given the known constraints of high rates of degradation of the generalist phage when cultured in minimal media. To do so, we imposed a cost of generalism not as burst size (or attachment rate), but instead as an increased intrinsic mortality rate for the generalist phage, while keeping the mortality rate the same for the three other species.

We observed that, when fitness cost was modeled as intrinsic mortality, our qualitative results matched those when fitness cost was measured as burst size or attachment rate (Figure 6; for burst size, see Figure 3A; for attachment rate, see Supplemental Figure 1A). In this case, the generalist phage could be maintained on competing prey even as its intrinsic mortality rate increased; competition with the specialist phage increased its rate of loss, but was insufficient to drive the generalist extinct across a wide range of parameters (Figure 6). However, if the intrinsic mortality rate of the generalist exceeds 5.7 times that of the specialist, it will be lost when bacteria compete. In contrast, comparable intrinsic mortality rates resulted in the early loss of the generalist when prey were mutualistic (when the intrinsic mortality rate of the generalist exceeds 2.6 times that of the specialist), with the specialist able to persist (Figure 6). However, it is worth noting that, because degradation rates *in vitro* are impacted by the physiological accessibility of the available prey and the differential abilities of the two phage to replicate on starved cells, our model is not currently designed to capture changes in intrinsic mortality rate as a function of prey density or physiological state. It is thus unlikely to perfectly replicate our experimental results. Regardless, in its current form, it reinforces our previous qualitative modeling predictions and our *in vitro* findings that suggest that competition between prey favors generalist predators across a wider range of fitness trade-offs than mutualism between prey.

**Figure 6.**
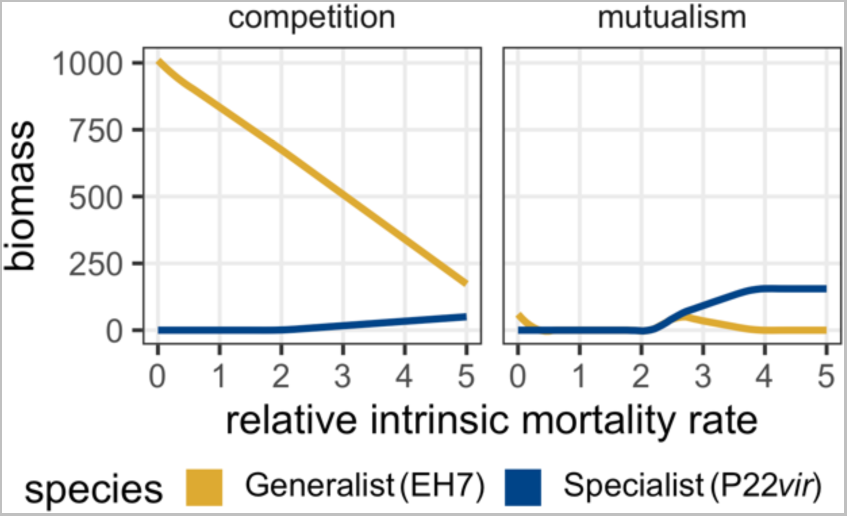
When a cost of generalism is modeled as intrinsic mortality, qualitative patterns of ecological selection on predator specificity match findings when cost of generalism is modeled as burst size (Figure 3) or attachment rate (Supplemental Figure 1). As the intrinsic mortality rate of the generalist phage increases, it maintains its advantage longer when prey compete and is driven extinct at an intermediate cost when prey are mutualistic. These qualitative results align with previous modeling findings when cost of generalism is imposed as burst size or attachment rate.

## Discussion

We aimed to determine whether ecological interactions between bacterial prey species impacted the abundance of phage with different prey specificity. We developed a simple four-species phenomenological model composed of two interacting bacterial species, a specialist phage, and a generalist phage. Using this chemostatic model, we found that specialist phage were favored when prey are mutualistic, while a generalist diet breadth was favored when prey compete. These results were robust to initial conditions across a range of parameter values, suggesting that they may be both ecologically and evolutionarily informative. We found that our modeling predictions were well-matched by the outcome of batch culture experimental phage competition assays, but that biological and environmental details not considered in our model contributed to these results. *In vitro*, phage degradation and inability to infect inactive bacteria drove ecological outcomes for phage. Our results highlight that ecological interactions between prey can alter predator specificity through a range of mechanisms.

Our modeling results suggest that interactions between bacterial prey impact the prevalence of phage specificity phenotypes. Experimental evolution has previously shown that the presence of different types of resources can select for generalism [9, 18, 73]. Both absolute and relative prey densities are relevant predictors of phage diet breadth [25, 26, 34, 35]. However, while much of the previous work done on diet breadth has assumed a constant relative abundance of available prey, our model upended that assumption by allowing relative prey abundances to vary as a function of prey ecology. Previous theoretical modeling has demonstrated that resource competition between prey species can select for expanded predator diet breadth even when trade-offs for generalism exist, although this result generally required the competitive dominance of the novel prey source [32, 33]. Our results align with these findings, underscoring that resource competition should favor a generalist strategy in most cases, even when a trade-off is present. Additionally, we expanded previous findings to include mutualistic interactions between prey, showing that a specialist predator strategy dominated assuming a minimal trade-off for generalism. Our model demonstrates that ecological interactions between prey species favor different predator diet strategies because switching from competition to mutualism changes relative prey abundances from being anti-correlated to being positively correlated. We anticipate that our modeling result will apply broadly to systems when interactions between prey generate correlations in their abundance.

The experimental results of our study largely align with our modeling predictions, although they also highlight two important aspects of our microbial system. First, *in vitro*, we did not observe a reproductive fitness cost of generalism on the shared prey species as we had anticipated in our model. Instead, we found that the trade-off manifested as an increased intrinsic mortality rate in minimal media and reduced replication rates on starved cells. Our results contribute to the body of work suggesting that pleiotropic costs are often context dependent [24, 36, 74, 75, 76, 77, 78, 79, 97]. Additionally, our experiments emphasize that in addition to altering population dynamics, interactions between bacteria can also impact phage diet breadth by altering prey physiology. Bacterial sensitivity to phage is rarely a binary trait and can change between physiological states due to differences in growth rate, metabolism, transcription and translational activity, and the availability of intracellular components [39, 40, 41]. Phage reproduction on slow-growing or stationary phase cells is often more difficult due to reduced cell size and lower densities of the receptors phage use to adsorb [25, 56]. Because microbial physiology is driven strongly by resource availability and interactions between bacterial species [39], our work suggests that ecological selection on phage specificity is likely impacted by how interactions between bacterial prey shift prey abundances and physiological states over time.

There are several limitations to the study we performed which impact the generality of the results. The two phage types tested differ significantly in traits in addition to diet breadth. They utilize different receptors for viral entry, with P22*vir* attaching to the O-antigen of *S. enterica*’s lipopolysaccharides (LPS) and EH7 using BtuB, a vitamin B12 uptake receptor. Previous work has demonstrated that BtuB expression is highly context dependent, while the LPS is constitutively expressed [80, 81, 82, 83, 84, 86, 87, 88, 89, 90, 91]. Additionally, our phages are vastly different sizes, with EH7 consisting of 110kb and significant similarity to phage strains with as many as 166 putative proteins, while P22*vir* is much smaller at 44kb and 74 proteins [53]. These differences are likely to accentuate the role of bacterial physiology in determining selective outcomes, as the intracellular demands for producing EH7 will be much higher and thus more limited by slow growth or low nutrient availability. Future work should test the competitive ability of phages that use the same or similar receptors and with greater stoichiometric similarity to disentangle these confounding factors. Additionally, we have not completed evolution experiments in this study to see if longer term selective outcomes can be well-predicted by ecological dynamics. However, in the context of microbiological engineering and biomedical applications, we expect that ecological selection will be a useful metric for predicting community outcomes.

Our results suggest numerous directions for future study. In the context of phage therapy, the performance of EH7 in minimal media emphasizes the necessity of testing how different environments affect phages and whether phage characteristics such as specificity tend to correlate with susceptibility to degradation [63]. These data also suggest that the ways bacteria modify their environments through alteration to local pH or vitamin concentrations may have consequences for the evolution and persistence of their viral predators. For example, human gut microbes often directly compete with their hosts for vitamin B12 [93]; the resultant availability of B12 in the human gut may alter the efficacy of BtuB-specific phages in phage therapy applications. Continued characterization of phage-bacteria interactions in the complex community contexts in which they are found will improve our ability to use phage for engineering and biomedical purposes. Finally, we note that the spatial structure of interacting bacterial species, as in a biofilm, will alter local prey availability in natural environments in ways that might complicate or invalidate the results we have presented here [36].

We took a simple modeling approach, paired with an ecological experiment, to gain insight into the role of prey ecology on the competitive ability of bacteriophage with different diet breadths. We found that, in both our model and *in vitro* experiments, prey interactions shape the prevalence of phage diet breadth phenotypes, though the mechanisms differ between our modeling approach and synthetic community. Management and design of microbial communities is contingent upon our ability to predict the evolutionary outcomes and higher order ecological effects of multitrophic interactions. Understanding the complex biotic factors driving ecological and evolutionary outcomes for bacteriophage is a critical step to effectively harnessing microbes for industrial and biomedical applications.

## Lab Acknowledgements

The authors thank X. Xiong, J. N. V. Martinson, J. Chacón, A. K. Shaw, M. Torstenson, C. Wojan, N. Narayanan, and D. Kim for helpful comments on the manuscript. The authors would also like to thank members of the Möbius and Nadell labs for input throughout the conceptualization and design of this project, as well as I. J. Molineux for providing our lytic P22*vir* strain. This work was supported by BBSRC via BBSRC-NSF/BIO grant BB/V011464/1 to W.M, and the National Science Foundation IOS-2017879 to C.D.N. and IOS-2019304 to W.R.H.

## Supplementary Analysis

### Degradation of the generalist phage in minimal media cannot be clearly attributed to any particular media component

We investigated the cause of phage degradation in our minimal media by incubating 10^3^ particles of EH7, P22*vir*, or a combination of both phage per well at 37°C with shaking at 432 rotations per minute in glucose minimal media for 48 hours. We found that EH7 degraded below the limit of detection within 24 hours whether incubated alone or with P22*vir*, half the time period of our standard phage competition assays (Supplemental Figure 2A). However, phage recovery was possible following the addition of cells in some cases (Figure 5B), suggesting that infectious phage particles remained. P22*vir* titer was unchanged over the course of 48 hours whether incubated alone or with EH7 (Supplemental Figure 2A, p = 1.0).

Given previous observations that environmental factors such as salinity [63, 64, 65, 66], pH [61, 62, 63], and metal presence can have significant impacts on phage titer [67, 68, 69], we attempted to identify the cause of degradation in our minimal media by removing one component at a time. We created minimal media without metals, without sulfur, without phosphorus, or some combination thereof. We incubated only 10^3^ EH7 particles per well at 37°C with shaking at 432 rotations per minute in each minimal media type for 48 hours. However, phage degraded below the limit of detection regardless of which media component was absent, suggesting that metals and osmolarity were not driving the reduction in titer (Supplemental Figure 2C). Additionally, because phage titer can be reduced as the result of adsorption of viral particles to plastic surfaces, we tested our phage by incubating it in LB [70, 71, 72]. We determined that adsorption of phage particles to the plastic was likely not driving the decrease in phage titer, given that EH7 density was unchanged following 48 hours of incubation in LB in the same 96-well plate where degradation in minimal media was observed (Supplemental Figure 2B, p = 0.00035). This was true despite the low starting phage density, which has been shown to increase the likelihood of rapid phage degradation due to adsorption to plastic [61, 70].

Finally, we investigated which component of LB was responsible for the preservation of phage titer by removing one component at a time. We repeated our previous experiments by incubating only 10^3^ EH7 particles per well at 37°C with shaking at 432 rotations per minute in each LB type for 48 hours. The titer of phage incubated in solutions containing tryptone were unchanged over the window of the 48 hour experiment (Supplementary Figure 2D). When incubated in a solution of yeast and salt, the density of EH7 particles was also not significantly changed (Supplementary Figure 2D). However, titer did decrease significantly relative to other conditions when phage were incubated in only yeast or only salt (Supplementary Figure 2D, yeast: p < 0.001 for all multiple comparisons, salt: p < 0.001 for all multiple comparisons). These results suggest that tryptone may play a role in the stability of EH7 titer in LB.

## Supplemental Figures

**Supplemental Figure 1.**
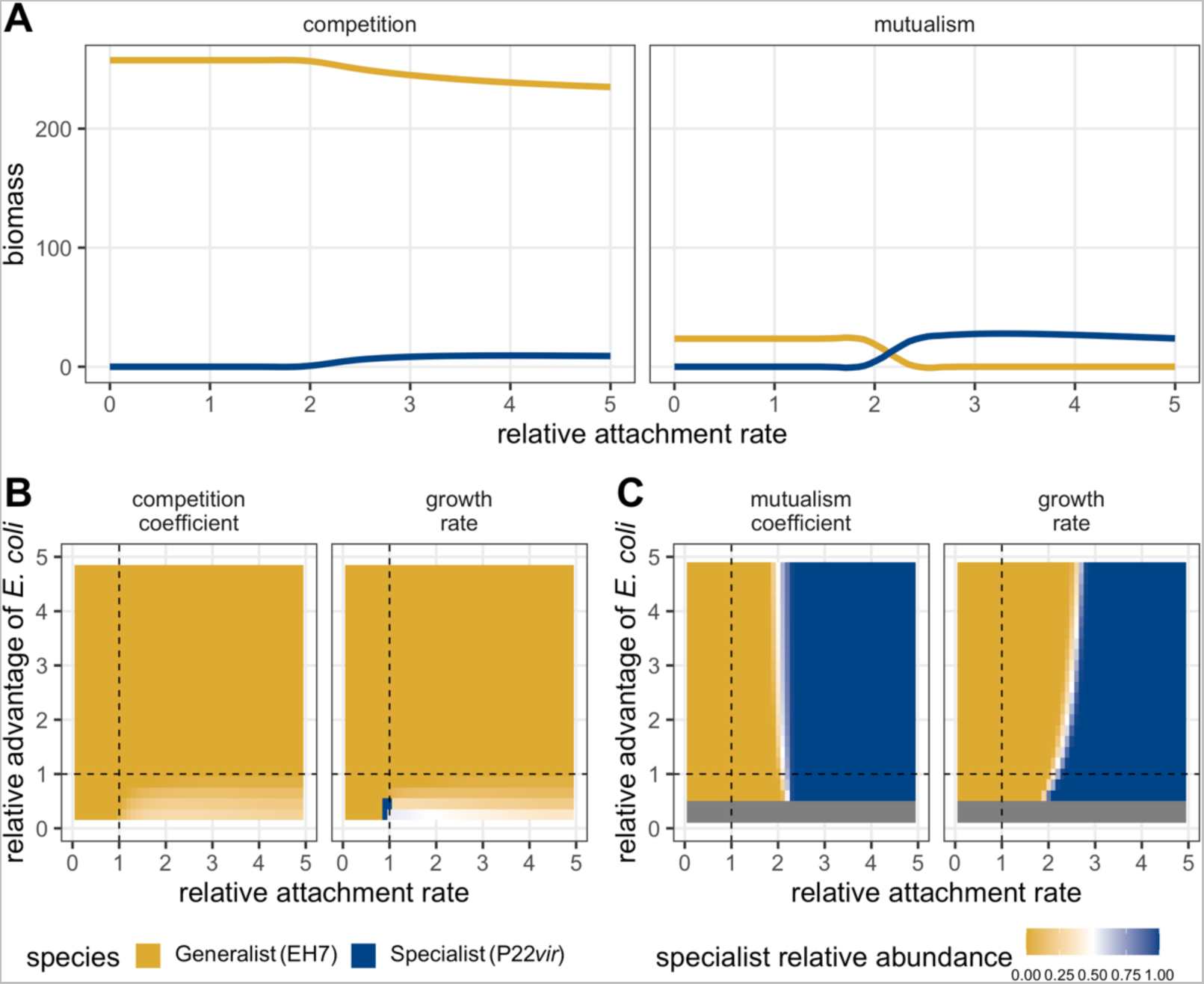
Numerically-simulated phage dynamics given a variety of parameter trade-offs demonstrate that prey interactions result in different patterns of predator abundance. **A:** The final density of each phage type as a function of bacterial interactions and increasing cost of generalism modeled as increasing specialist attachment rate. When prey are mutualistic, a cost of generalism exists that favors specialists (blue line) over generalists (yellow line). When prey compete, no such cost exists; this is true even as the specialist’s attachment rate increases well beyond the values displayed here. **B:** The relative abundance of the specialist phage on competing prey as a function of increasing cost of generalism and relative growth advantage of the alternative prey *E. coli*. Whether prey growth advantage is modeled through growth rate or competitive coefficients (beta), the generalist is favored (yellow) except in a small subset of cases where the alternative prey is competitively excluded. **C:** The relative abundance of the specialist phage on mutualistic prey as a function of increasing cost of generalism and relative growth advantage of the alternative prey *E. coli*. Whether prey growth advantage is modeled through growth rate or mutualistic benefit (alpha), a cost of generalism exists above which specialism is favored (blue). Note that there are benefit and growth rate values for *E. coli* below which the system cannot be supported, indicated by the grey bar.

**Supplemental Figure 2.**
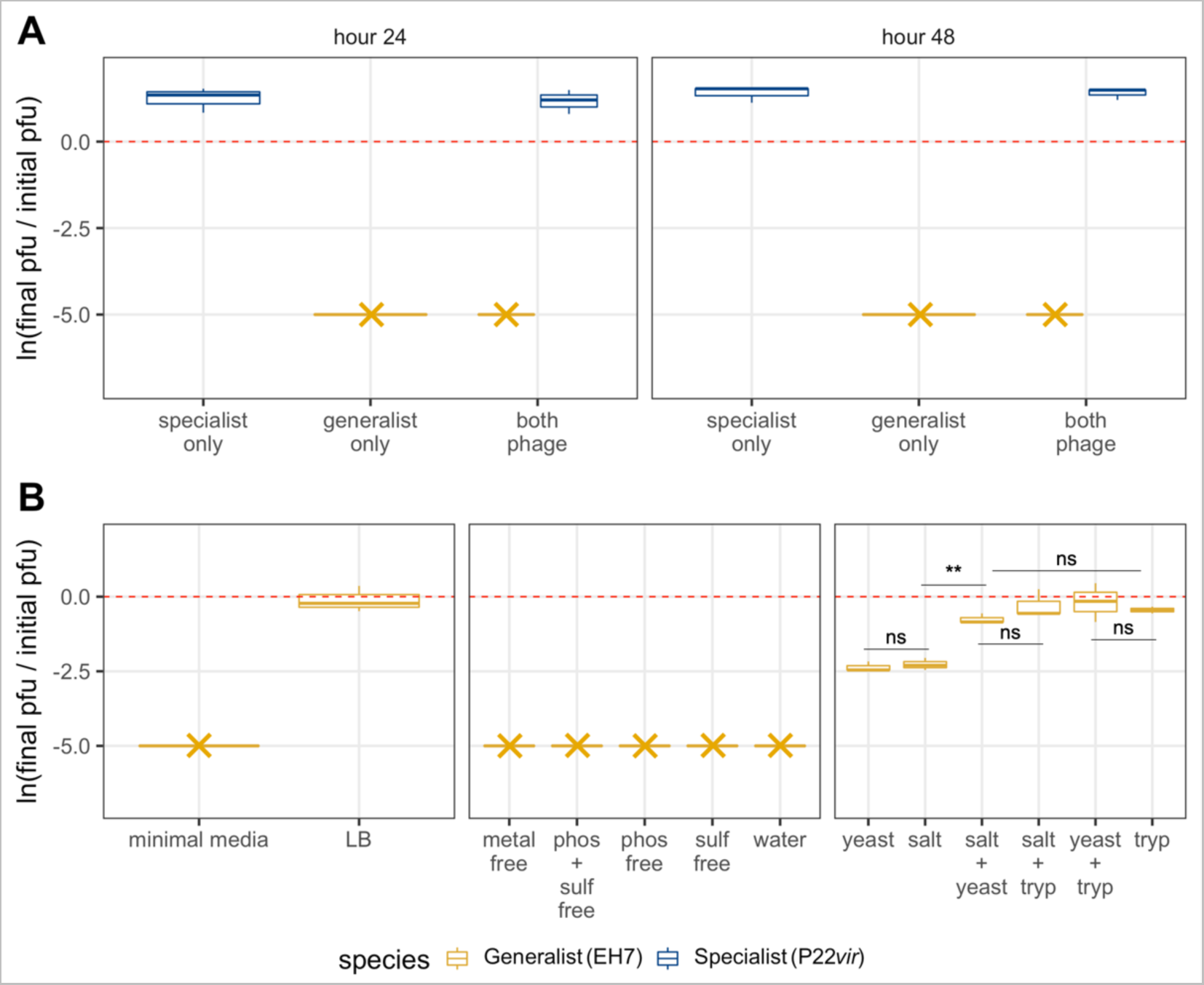
Generalist phage degradation differs in minimal media versus LB. **A:** EH7 is undetectable after 24 and 48 hours, as indicated by the yellow asterisks, when incubated in minimal media without cells. However, because replication is still possible when cells are added at 24 hours (see Figure 5B, where EH7 titer increases on competitive co-culture even when the addition of cells is delayed), these results suggest that phage are below the limit of detection but some infectious particles remain. Statistical significance was determined using a one-way ANOVA with Tukey’s HSD multiple comparison test. **B:** Change in EH7 titer across media conditions. In LB without cells, EH7 densities are unchanged over a 48 hour period, while the phage disappears below the limit of detection in minimal media (p = 0.00035), as indicated by the yellow asterisk. Statistical significance for the first facet was determined using a two-tailed t-test. EH7 degradation is not impacted when different components of the minimal media are removed - it is always undetectable after 48 hours, as indicated by the yellow asterisks. Regardless of what component is removed, the phage is undetectable after 48 hours. Change in EH7 titer does differ when different components of LB are removed. Phage is always detectable when at least one component of LB is present, while solutions containing tryptone or both yeast and salt best preserve phage titer over 48 hours of incubation (Supplemental Analysis 2). Statistical significance for the third facet was determined using a one-way ANOVA with Tukey’s HSD multiple comparison test. **Note:** in part B, facet panels indicate experiments were completed on different days. **For A and B:** The dotted red line indicates no change in titer from the start of the experiment to the end. Values greater than zero indicate an increase in titer, while values below zero indicate a decrease in titer.

## Supplemental Tables

**Supplemental Table 1.** Phage and bacterial strains. Strains used for experiments. See materials and methods for additional details.

| Strain ancestor | Antibiotic marker | Flourescent marker | Source | Relevant phenotype |
| --- | --- | --- | --- | --- |
| Salmonella enterica LT2 metA* metJ* | kanR | YFP | Harcombe 2010 | Methionine hypersecreter |
| E. coli K-12 BW25113 ΔmetB | kanR | CFP | Harcombe 2010 | Methionine auxotroph |
| E. coli K-12 BW25113 ΔtrxA | kanR | NA | Baba et al. 2006 | Sensitive to EH7, resistant to P22vir |
| S. enterica serovar Typhimurium NCTC 74 ΔbtuB | NA | NA | S. Bowden | Sensitive to P22vir, resistant to EH7 |
| P22vir | NA | NA | I. J. Molineaux | S. enterica-specific phage |
| EH7 | NA | NA | E. Hansen | Generalist phage |

**Supplemental Table 2.**
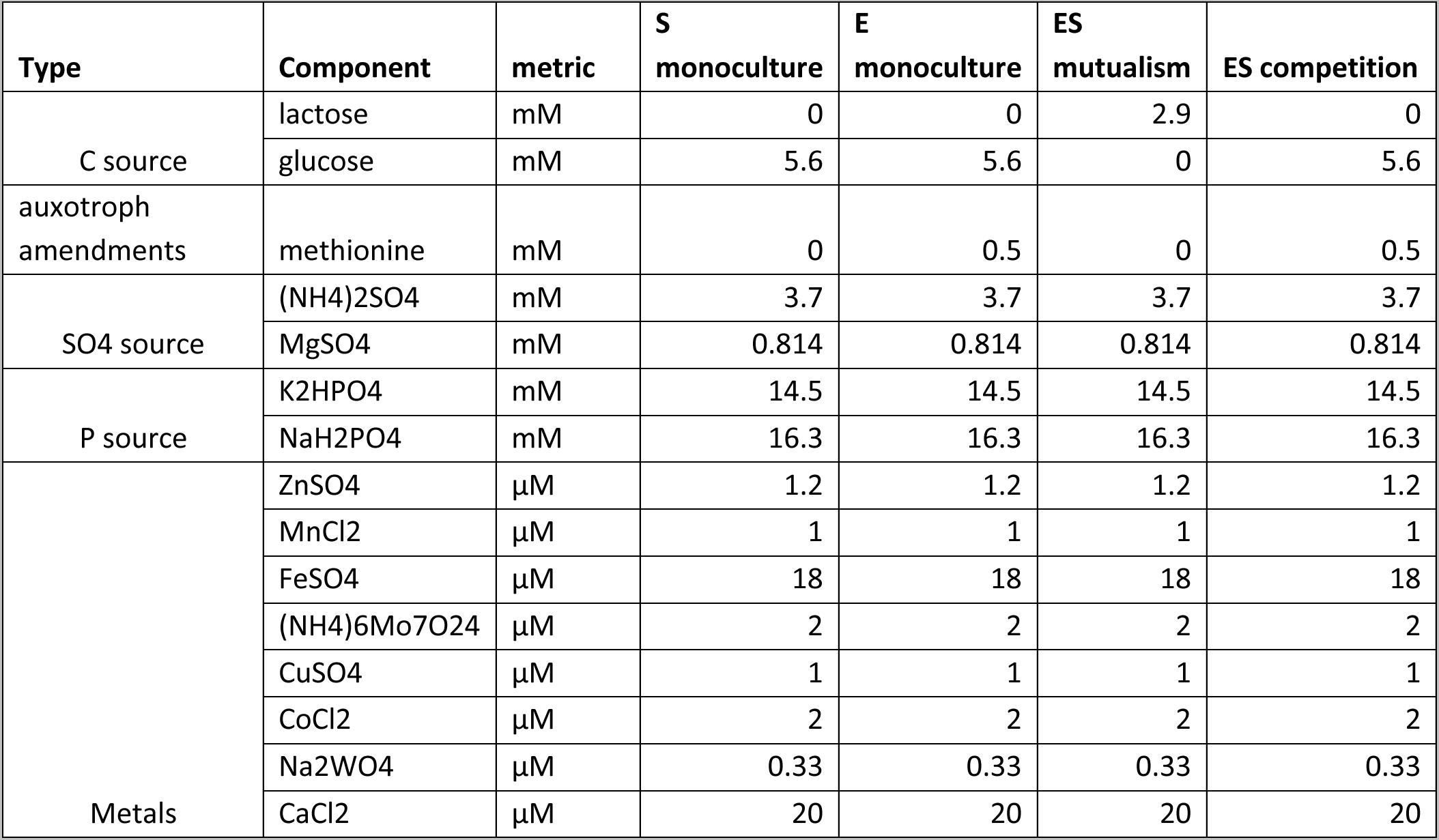
Hypho minimal media composition. Media composition for mutualistic and competitive communities, as well as bacterial monocultures. Note that phage degradation assays on starved cells were completed in ES mutualism media.

| Type | Component | metric | S<br>monoculture | E<br>monoculture | ES<br>mutualism | ES competition |
| --- | --- | --- | --- | --- | --- | --- |
| C source | lactose | mM | 0 | 0 | 2.9 | 0 |
|  | glucose | mM | 5.6 | 5.6 | 0 | 5.6 |
| auxotroph<br>amendments | methionine | mM | 0 | 0.5 | 0 | 0.5 |
| SO<sub>4</sub> source | (NH<sub>4</sub>)<sub>2</sub>SO<sub>4</sub> | mM | 3.7 | 3.7 | 3.7 | 3.7 |
|  | MgSO<sub>4</sub> | mM | 0.814 | 0.814 | 0.814 | 0.814 |
| P source | K<sub>2</sub>HPO<sub>4</sub> | mM | 14.5 | 14.5 | 14.5 | 14.5 |
|  | NaH<sub>2</sub>PO<sub>4</sub> | mM | 16.3 | 16.3 | 16.3 | 16.3 |
| Metals | ZnSO<sub>4</sub> | μM | 1.2 | 1.2 | 1.2 | 1.2 |
|  | MnCl<sub>2</sub> | μM | 1 | 1 | 1 | 1 |
|  | FeSO<sub>4</sub> | μM | 18 | 18 | 18 | 18 |
|  | (NH<sub>4</sub>)<sub>6</sub>Mo<sub>7</sub>O<sub>24</sub> | μM | 2 | 2 | 2 | 2 |
|  | CuSO<sub>4</sub> | μM | 1 | 1 | 1 | 1 |
|  | CoCl<sub>2</sub> | μM | 2 | 2 | 2 | 2 |
|  | Na<sub>2</sub>WO<sub>4</sub> | μM | 0.33 | 0.33 | 0.33 | 0.33 |
|  | CaCl<sub>2</sub> | μM | 20 | 20 | 20 | 20 |

**Supplemental Table 3.** Sobol’ sensitivity analysis and Morris screening parameter ranges. Minimum and maximum uniform distribution values used for each ODE parameter in Morris screening and Sobol’ sensitivity analyses.

| Parameter | Minimum | Maximum |
| --- | --- | --- |
| $\alpha_{j,i}$ | 0.1 | 2.5 |
| $\beta_{j,i}$ | 0.1 | 2.5 |
| $\mu_i$ | 0.1 | 2.5 |
| $\gamma_{i,x}$ | 15 | 65 |
| $\zeta_{i,x}$ | 0.0009 | 0.01 |
| $\delta_i$ | 0.0009 | 0.1 |
| $\kappa_i$ | 0.1 | 10 |
| $R$ | 0 | 5 |

**Supplemental Table 6.** Breseq predictions of point mutations responsible for repressing lysogeny in P22*vir*. Breseq predictions of point mutations in P22*vir* strain used for all experiments relative to the ancestral, lysogenic version of P22 (GenBank accession NC_002371.2). Sequencing of the lab phage strain was completed by the Microbial Genome Sequencing Center (https://www.seqcenter.com/).

| Predicted mutations |  |  |  |  |  |
| --- | --- | --- | --- | --- | --- |
| position | mutation | freq | annotation | gene | description |
| 30,899 | G→T | 100% | intergenic (-240/+114) | 24 ← / ← c2 | unknown/prophage repressor |
| 31,700 | C→G | 100% | intergenic (-37/-44) | c2 ← / → cro | prophage repressor/repressor |
| 31,716 | A→G | 100% | intergenic (-53/-28) | c2 ← / → cro | prophage repressor/repressor |

